# No evidence of “social immune memory” in Acorn ants (*Temnothoax nylanderi*)

**DOI:** 10.1101/2023.10.20.563152

**Authors:** Jaya Robinson, Christopher D. Pull

**Author notes:** Corresponding author: Christopher D. Pull, 11a Mansfield Road, Oxford, OX1 3SZ, UK.

## Abstract

Social organisms are predicted to experience higher rates of infectious disease, as the likelihood of transmission increases with higher spatial and temporal proximity of potential hosts. In social insects, this risk is counteracted through a social immune system, where altruistic and collective anti-pathogen behaviours reduce or eliminate the impact of disease at the colony, or ‘superorganism’, level. Superorganismal immune strategies are often analogous to immune responses in metazoan organisms, suggesting that disease may have played a crucial role in shaping multiple major evolutionary transitions. In this study, we investigated whether acorn ants use prior experience with a pathogen to perform more efficient social immunity behaviours upon pathogen reencounter, a potential parallel to metazoan immune memory that we term ‘social immune memory’. Using *Temnothorax nylanderi* ants, we created experience with the fungal pathogen *Metarhizium brunneum* both individually and in groups, to investigate the effect of prior experience when presented with the same pathogen. We further examined the effect of prior pathogen experience when groups of ants differ in their pathogen-experience levels. We found no evidence for social immune memory in our experiments, suggesting positive results in previous studies may be due to sublethal infections, rather than cognitive effects. Alternatively, ants may employ different immune defence strategies depending on context or species, with potential differences in immune system flexibility across major evolutionary scales.

## INTRODUCTION

Social animals face a significant risk of infection from conspecifics, that is thought to be amplified by living in groups and interacting more frequently with others (Ezenwa et al., 2016; Rifkin et al., 2012). Indeed, epidemiological SIR modelling demonstrates that the infectious potential (R_0_) of many pathogens is increased by factors typical of animal sociality (Schmid-Hempel, 2017). Despite the predicted increase in infection risk, many animals nevertheless adopt social lifestyles, often mitigating this risk through behavioural adaptations. For example, many social organisms avoid conspecifics showing signs of infection, including humans (Curtis, 2014), fish (Stephenson et al., 2018), and social insects (Drum & Rothenbuhler, 2015). Where groups consist of closely related individuals, there is the potential for kin-selected, altruistic disease-mitigating behaviour to emerge, which is suggested to occur in both humans (Shakhar & Shakhar, 2015) and mandrills (Poirotte & Charpentier, 2020).

Social insect colonies with distinct castes (ants and termites, and social bees and wasps) are a remarkable example of socially living animals. Typically consisting of a queen and her daughters, these colonies exhibit reproductive division of labour, with workers foregoing reproduction to instead help raise the queen’s brood, who is herself reliant upon her workers for survival and brood care (Boomsma & Gawne, 2018). Where castes are differentiated, this interdependence is fixed, such that selection acts upon the social insect colony itself, a colonial superorganism, with constituent parts that aim to maximise the superorganism’s fitness (Boomsma & Gawne, 2018); the emergence of superorganisms is thus considered one of the major evolutionary transitions (West et al., 2015). Social insects behave cooperatively to counteract the threat of disease, forming a social immune system that reduces the risk of infection at the level of the colony (Cremer et al., 2018). These superorganismal immune defences often demonstrate remarkable similarities to the physiological immune systems of many multicellular organisms (Cremer & Sixt, 2009). Such parallel evolution, occurring at different levels of biological organisation, suggests that similar selection pressures may have been operating across the major evolutionary transitions. Indeed, it has even been suggested that the presence of an immune system which operates at the new, higher level of biological organisation may be requisite to enable major evolutionary transitions in the first place (Meunier, 2015; Pradeu, 2013; Pull & McMahon, 2020).

One largely unexplored parallel between organismal and superorganismal immune strategies is that of immune memory. This is a key component of many metazoan immune systems, where pathogen-specific cellular immune mechanisms are enacted faster and to a stronger extent in individuals with prior experience of a pathogen (Janeway et al., 2005). The typically sedentary, long-lived lifestyle of ant colonies is expected to result in recurrent pathogenic infections, and experience-enhanced social immunity could benefit superorganisms through both reduced sanitary care error and reduced pathogen presence. Indeed, ants have previously been shown to integrate prior experience to improve both thermal (Weidenmüller et al., 2009) and foraging behaviours (Ravary et al., 2007). We therefore predict that ants may demonstrate behaviours producing, at the colony level, anti-pathogen mechanisms analogous to organismal immune memory, which we term ‘social immune memory’.

Previous work has demonstrated that ants exposed to pathogen-contaminated nestmates show higher levels of allogrooming, a social immune mechanism where insects remove fungal spores from colony members’ bodies (Reber et al., 2011; Walker & Hughes, 2009; Westhus et al., 2014). However, these findings may result from sub-lethal infections acquired during the first pathogenic exposure that are known to cause changes in social immunity behaviour (Konrad et al., 2018). In addition to immune priming, sublethal infections can lead to reduced interaction with the brood (Bos et al., 2012), through physiological effects rather than any effects of experience that would be dependent on learning and memory (Konrad et al., 2018). Additionally, while some studies have demonstrated a behavioural effect of pathogen experience, it has not been sufficiently examined if these effects are dependent upon secondary encounter with the pathogen (Reber et al., 2011; Westhus et al., 2014). Where this has been investigated, behavioural changes have only been observed at the group level, and neither individual ant behaviour, nor scenarios of mixed experience levels in groups, have been investigated (Goes et al., 2023; Walker & Hughes, 2009).

In this study, we aim to address this gap in our understanding of social immunity behaviour. We performed experiments on *Temnothorax nylanderi* ants which, due to workers’ long lifespan and solo foraging behaviours that are dependent on individual experience, are an ideal model system to investigate the role of individual experience on social immunity. We conducted experiments on individuals by repeatedly removing 48 ants from their colony and exposing them to larvae coated in either a control solution or fungal conidiospores that were UV-irradiated to prevent infection of the ants. Following this training period, we introduced these experienced ants to fungal-contaminated larvae and examined the grooming behaviour of pathogen-experienced and the previously pathogen-naïve controls. We also performed a group experiment, housing ants together for a week and creating pathogen experience by introducing fungal-contaminated larvae daily, again investigating ant behaviour when contaminated larvae were introduced. We further created mixed groups of pathogen-experienced ants with pathogen-naïve nestmates, analysing behaviour when presented with pathogen-exposed brood. By using both a live (infectious) and UV-irradiated (non-infectious) pathogen, our aim was to disentangle effects resulting from physiological processes from that of prior behavioural experience, in both our group experience experiment and naïve nestmate experiment.

## MATERIALS & METHODS

### Ant colonies

We used *Temnothorax nylanderi* ants for all experiments. We collected mature colonies in October 2022 from mixed deciduous forest within the university parkland of Royal Holloway University of London, where this species is locally abundant (Pontin, 2005). We housed colonies in a controlled temperature room at the University of Oxford (21°C with a 12:12hr light/dark cycle), supplying each with a*d libitum* 30% sucrose solution, and fruit flies (*Drosophila melanogaster*) every week. All colonies used in experiments were monogynous and ranged in size from 48-358 workers. When selecting larvae for our experiments, we chose the largest larvae in each colony (all late-instar larvae).

To identify ants throughout experiments, we painted workers with a paint-dot ID system. Despite widespread use of this approach (Pratt et al., 2002; Walker & Hughes, 2009), we found paint marks were removed through self-and allogrooming in many cases. We achieved best paint retention by removing and painting all workers, applying enamel paints (Testors Pactra Racing Finish, USA or Mr Color) with a fine nylon wire, and then placing the painted ants into entirely new nests that we had already transferred their brood into. We painted each ant with four spots, using seven potential colours, with each ant receiving a unique colour code that was robust to the loss of one paint spot (Burchill & Pavlic, 2019). Using this approach, we were able to produce enough painted ants that were uniquely identifiable for experiments and lost only three replicates due to paint loss.

### Fungal pathogen

We used the entomopathogenic fungus *Metarhizium brunneum* (isolated from an infected *Lasius niger* queen; Pull & Cremer 2017). This is an opportunistic pathogen that infects a wide range of insects, including ants, by conidiospores attaching to the body surface, before germinating and penetrating the cuticle to cause infection (Castrillo et al., 2005). Prior to infection, conidiospores can be removed from ants’ body surfaces through self-and allogrooming. Conidiospores were harvested and suspended in sterile (autoclaved) 0.05% Triton X-100 (Sigma-Aldrich Co., USA).

To distinguish behaviours resulting from pathogen experience from those caused physiologically by sub-lethal infection, we created UV-inactivated conidiospores to simulate pathogen exposure but without causing infections in our grooming ants (Braga et al., 2001). UV-inactivated conidiospores were produced by placing 6ml of 10^7^ conidiospore solution in a 90mm petri dish (Thermo Fisher Scientific, Inc.) and exposing the suspension to 254 nm UV light for 1 hour (Vilber Bio-Link 254; 230 W). To check that UV-irradiation was successful, samples were incubated on sabouraud dextrose agar plates at 25°C for one week before use, and for the duration of the experiment; no colony forming units were ever present on plates nor any production of hyphae from spores when viewed under microscope.

To contaminate larvae with conidia, we used sterile (ethanol-wiped) soft forceps to roll single larva in a 0.3μl droplet of either 1×10^8^ conidiospore suspension or a control solution of autoclaved 0.05% Triton X-100, on sterile (ethanol-wiped) parafilm, leaving larvae to air-dry for 10-20 mins.

### Individual experiments

To investigate the effect of prior fungal experience on individual ant behaviour, we repeatedly exposed ants to larvae coated in either a UV-irradiated pathogen or a control solution (‘training phase’; Figure 1), and then investigated the effect of differential experience levels when all ants (pathogen-experienced and control) were exposed to fungal-contaminated larvae (‘test phase’; Figure 1). We placed workers individually into arenas (30mm petri dishes [Thermo Fisher Scientific, Inc.], plaster of Paris bases), allowing them to acclimate for 10-20 min. We then placed a single larva in the arena that had been coated in either UV-sterilised fungus (pathogen-experienced ants) or 0.05% Triton X (control ants). Ants were left with these larvae for 2 h, after which workers were returned to their colonies. We repeated this exposure period every two days, for a total of five times per worker. Two days after this ‘training phase’, we placed both pathogen-experienced and control-experienced ants with larvae exposed to UV-sterilised fungus, and video recorded their behaviour during this ‘test phase’ (Figure 1). We used a total of 12 colonies where all ants had been painted. To avoid the spread of fungal conidia from pathogen-experienced to control ants, we assigned all experimental ants from a colony to one of these two treatments. We selected four workers per colony, choosing those observed caring for the brood (n = 24 control & 24 pathogen-experienced; 48 total). To examine grooming efficacy, we determined the number of conidiospores remaining on the larvae after being groomed in the arenas. To determine if ants improved in grooming efficiency over time, we compared the number of remaining conidiospores for all ants assigned to the UV pathogen experience treatment between their first and last exposure period. To measure the effect of prior pathogen experience, the number of remaining conidiospores were compared between larvae groomed by ants assigned to either experience treatment during the test phase (Figure 1). Following exposure periods, we placed larvae individually in 500μl microtubes (Kartell Spa Noviglio, USA) containing 5μl Triton X-100. We vortexed these at a high speed for 10 mins (S0200-Vortex Mixer, Labnet International Inc, USA), placing 2μl of this solution into a haemocytometer. We then counted the number of conidiospores in both counting chambers for each larva. We repeated this for 20 control larvae, which received identical treatments but were not groomed. We compared the number of remaining conidiospores between larvae groomed by pathogen-experienced and control ants, and conidiospore numbers for larvae groomed by the same ant during its first and last exposure period.

**Figure 1.**
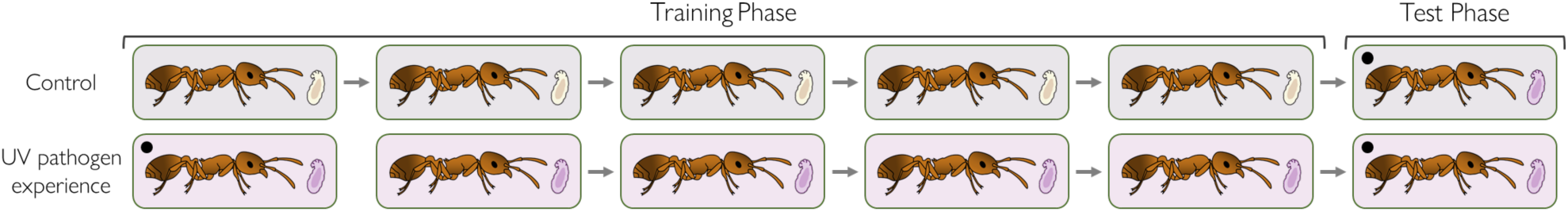
Experimental procedure for individual experiments. We removed workers from original colonies, placing them in arenas (boxes) with larvae for 2h. We repeated this five times to form the training phase, with two-day intervals between exposure periods (arrows). During the training phase, control ants were given larvae coated in Triton X-100 (clear larvae), and UV-pathogen-experience ants were given larvae coated in UV-irradiated *M. brunneum* (purple larvae). Two days after the training phase, all ants were exposed to larvae coated in UV-irradiated *M. brunneum*. Boxes are coloured by treatment group. Black circles denote larvae that were subsequently examined to determine the number of conidiospores remaining on the surface. We repeated this procedure for four ants from each of 12 colonies (n = 24 control & 24 pathogen-experienced; 48 total)

### Group experiments

To investigate the effect of pathogen experience on individual behaviour in groups, we performed experiments on groups of five workers that experienced repeated exposure to treated larvae, following a similar protocol to above. We housed five workers from the same colony together in arenas (30mm petri dishes, plaster of Paris bases), supplied with 30% sugar water *ad libitum* (plastic dish with sugar water-soaked cotton ball). We repeated this to form five groups, for each of five colonies (n = 125 ants). To identify individual behaviour in these groups, we painted the thorax and abdomen of workers with a single colour (either green, yellow, orange, white, or blue) immediately before placing them in arenas. We allowed these groups to acclimate for one day, repainting a small number of individuals who had removed their paint spots. We then introduced three larvae into each of these groups: for each colony, one group of workers received larvae coated in live (infectious) fungal pathogen, one group received larvae coated in UV-sterilised (non-infectious) fungal pathogen, and three groups received larvae coated in Triton X-100, as our experimental design required multiple control groups (Day 1-5; Figure 2). We left larvae in arenas for 6h before discarding them. We repeated this procedure daily for five days, checking nests and removing dead workers without replacement (< 7 total across treatments) 2h before larval introduction. On the 6^th^ day, we introduced three larvae coated in live fungal conidiospores to both the live-pathogen-experience and a control group; and three larvae coated in UV-irradiated conidiospores to the UV-pathogen-experience group, as well as a control group (Day 6; Figure 2). We also introduced three larvae coated in Triton X-100 to the third control group for use as controls in the subsequent naïve nestmate grouping experiment. To record potential differences in behaviour between pathogen-experience and control groups on this day, we conducted behavioural scan samples every 4 minutes for 1.5h after larval introduction, recording the behaviour of each ant (behavioural categories included: *brood grooming, self-grooming, allogrooming, brood carrying, sat with brood, venom gland grooming, walking,* and *inactivity*). Two researchers carried out scan sampling, one to observe the ants and one to record the behaviour, and the observer was blind to the treatment groups.

**Figure 2.**
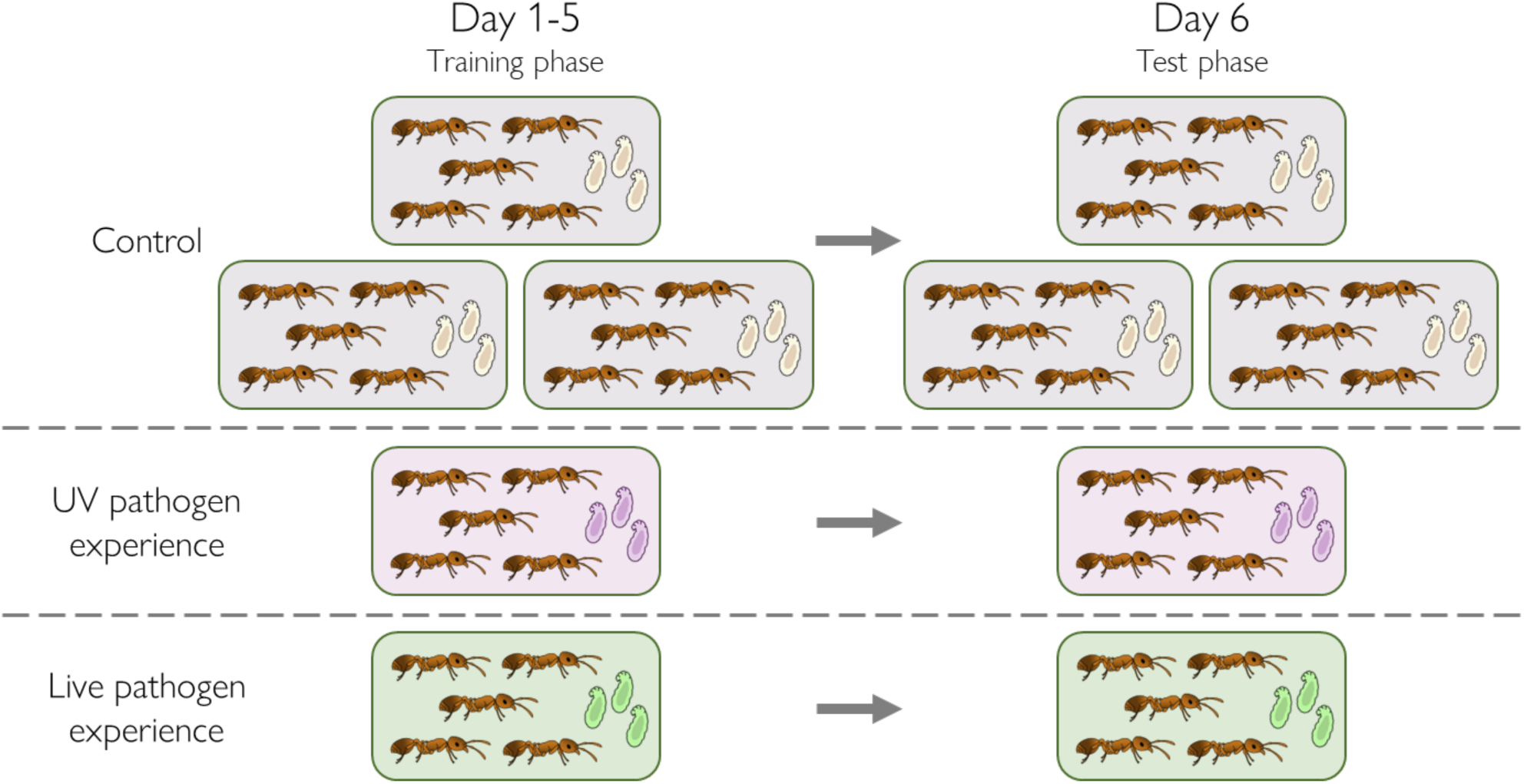
Experimental procedure for group experiments. We created five groups of five workers, which we housed in arenas (boxes). For the first five days (days 1-5, training phase) we introduced three larvae to the groups for a duration of 6h, after which larvae were removed from arenas. Three groups were assigned to the control treatment, and received larvae coated in Triton X-100 (clear larvae) during this training phase; one group was assigned the UV-pathogen-experience treatment, and received larvae coated in UV-irradiated *M. brunneum* (purple larvae); and one group was assigned to the live-pathogen-experience treatment, and received larvae coated in live (infectious) *M. brunneum* (green larvae) during the training phase. Following this training phase (day 6, test phase), groups were presented with contaminated larvae for 1.5h, during which we performed behavioural scan samples. On this day, we introduced larvae coated in UV-irradiated *M. brunneum* to one of the control groups and the UV-pathogen-experience group, and we introduced larvae coated in live (infectious) *M. brunneum* to a second control group and the live-pathogen-experience group. We introduced larvae coated in Triton X-100 to the third control group, which were used in the naïve nestmate grouping experiment. Boxes are coloured by treatment group. We carried out this procedure for each of five experimental ant colonies (n = 100)

### Naive nestmate experiments

To explore behavioural responses when workers differ in pathogen experience, we investigated the individual behaviour of ants in mixed groups of pathogen-experienced and pathogen-naïve individuals. For this, we used pathogen-experienced ants from the previous group experiment, and ants which had undergone the same experimental procedure, at the same time, but had no prior experience with the pathogen (see above). Following the final day of the group experiment (Day 6; Figure 3), we placed one of these workers in a new arena with two pathogen-naïve nestmates that had been acclimating for one day (arenas supplied with 30% sucrose water). To distinguish experienced and naïve ants, we painted the thorax and abdomen of naïve nestmates with orange dots. For each of the five colonies, we created two groups of (i) one UV-pathogen-experienced ant and two naïve nestmates (UV experience), (ii) one control ant and two naïve nestmates (UV control), (iii) one live-pathogen-experienced ant and two naïve nestmates (live experience), and (iv) two more groups of one control ant and two naïve nestmates (live control) (n = 120 ants; Figure 3). We then allowed ants to acclimate for 30 minutes and then introduced one contaminated larva to each arena. To UV experience and UV control groups, we introduced larvae coated in UV-sterilised conidiospores, and added larvae coated in live (infectious) conidiospores to live experience and live control groups (Figure 3). These exposure periods lasted 1.5h, and we performed scan-sampling (as above) every 4 minutes for 1.5h using the same behavioural categories as before.

**Figure 3.**
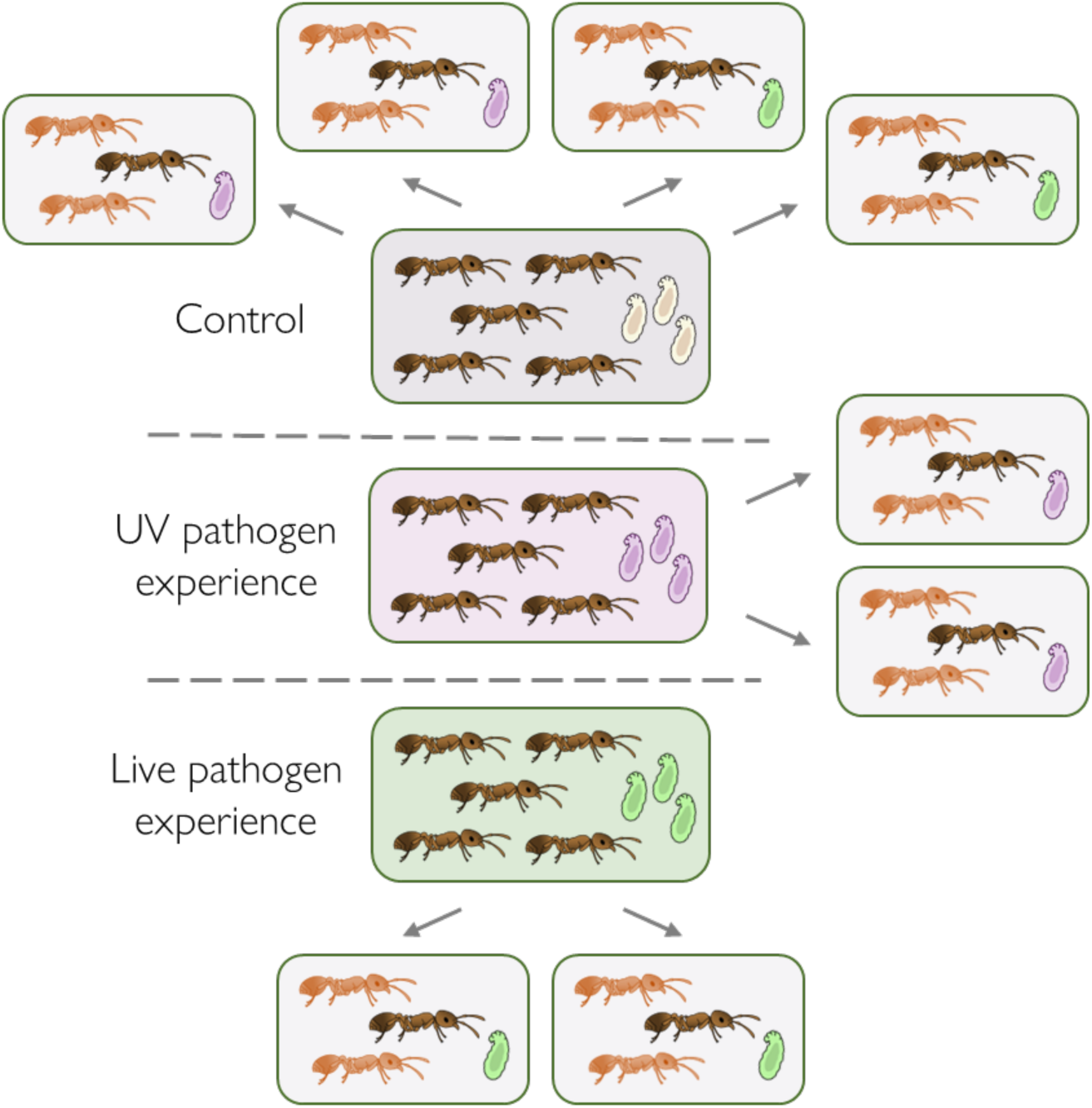
Experimental procedure for naïve nestmate experiments. Using ants from the group experience experiment (grey ants), we created groups with different levels of pathogen experience by housing these with pathogen-naïve nestmates (orange ants) from their original colonies. Following the group experiment, we placed four ants from the control group which had only experienced larvae coated in Triton X-100 (dark grey box) individually in arenas (light grey boxes) each with 2 nestmates. Similarly, from both the UV-pathogen-experience (purple box) and live-pathogen-experience (green box) groups, we removed two ants and placed them individually with two pathogen-naïve nestmates. We then introduced 1 larva to each of these arenas: two control groups and both groups with UV-pathogen-experienced ants received larvae coated in UV-irradiated *M. brunneum* (purple larvae); and the remaining two control groups and both groups with live-pathogen-experienced ants received larvae coated in live (infectious) *M. brunneum* (green larvae)

**Figure 4.**
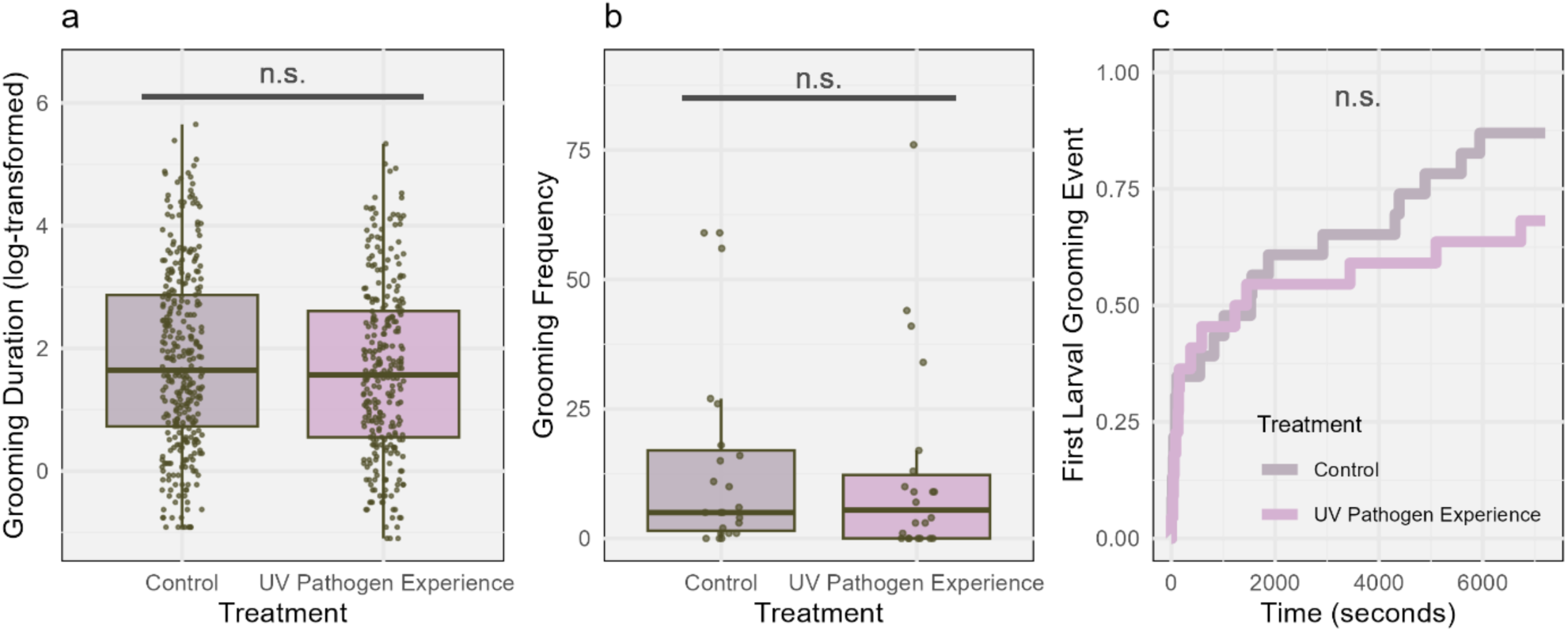
Effects of individual experience on grooming behaviour. (A) Duration (seconds) of each grooming event for control ants (grey) and UV-pathogen-experienced ants (purple) (all data points displayed, values have been log-transformed for visualisation purposes, jitter = 0 vertically, 0.1 horizontally); (B) Frequency of grooming events during the observation period for control ants (grey) and UV-pathogen-experienced ants (purple) (all data points displayed, jitter = 0 vertically, 0.1 horizontally); (C) Cumulative proportion of control ants (grey) and UV-pathogen-experienced ants (purple) grooming larvae during the experimental period. Boxplot features: centre line = median, box limits = 1^st^ and 3^rd^ quartiles (Q1 to Q3), whiskers = from Q1 to smallest value, at most –1.5 × IQR (inter-quartile range; Q3-Q1); and from Q3 to largest value, at most +1.5 × IQR. Letters above boxplots depict results of pairwise significance testing between groups, n.s. = non-significant

### Video analysis

We recorded exposure periods using a Raspberry Pi camera system (Raspberry Pi 4 Model B V1.2), by mounting cameras on a clamp stand and placing arenas directly below. To record individual experiments, we used four cameras, each recording six arenas, meaning that all exposure periods occurred simultaneously. We recorded our group and naïve nestmate experiments in a similar manner, performing half the exposure periods in the morning, and immediately repeating the procedure for remaining arenas in the afternoon. However, we declined to analyse these video recordings as our scan sampling data was sufficient to test our hypotheses. We analysed recordings of individual experiments using BORIS software (v8.11.4; Friard & Gamba, 2016) to manually code behaviours from our videos. We recorded the time that larvae were placed in arenas, and the start and end of larval grooming periods for 2h after introduction. We blinded this process by ensuring treatment configuration was unknown during analysis.

### Statistical analyses

We performed all statistical analyses in R (v4.2.3; R Core Team, 2021), using ‘lme4’ and ‘coxphf’ packages to model our data (Bates et al., 2014; Heinze et al., 2020), and the ‘ggplot2’ package to plot all graphs (Wickham, 2009). We checked the necessary assumptions of all tests by, where appropriate, viewing histograms of data, plotting the distribution of model residuals, checking for non-proportional hazards, testing for unequal variances, testing for the presence of multicollinearity, testing for over-dispersion, and assessing models for instability and influential observations. Where relevant, we fitted an observation-level random intercept effect to address overdispersion (Harrison, 2014). All models were compared to a null (intercept-only) model that had the same random effect structure using a likelihood ratio (LR) test (Bolker et al., 2009) and, where necessary, a reduced model to test the impact of multiple fixed effects. We included a random intercept effect for colony in all models as ants from the same colony are non-independent due to their shared genetic background and experience and, where appropriate, included a random intercept effect for time of day that experimental sessions were carried out (morning or afternoon); for the Cox proportional hazards model we used the inbuilt frailty function with a gamma distribution to the incorporate random effect (Austin, 2017). Where models had singular fit, we removed random effects with negligible variance (σ^2^ < 1×10^-5^) to simplify the models. Where we performed multiple tests on one experiment, we used the Benjamini-Hochberg procedure to control the false discovery rate (α=0.05; Benjamini & Hochberg, 1995; García, 2004). Where we found significant fixed effects, we performed post-hoc tests using the ‘multcomp’ package to determine which groups were significantly different, using Tukey’s HSD method with the Benjamini-Hochberg procedure to control for multiple comparisons (Hothorn et al., 2008).

### Individual experiments

To analyse our individual experiments, we calculated the time until the first instance of larval grooming (latency) and the number of grooming events (frequency) during the 2-hour observation period for each ant, as well as the duration of each grooming event (duration). Due to paint spot removal, 3 out of 48 ants were unidentifiable on the final day, therefore our analyses of individual experience used data from 45 ants (22 fungal-experienced, 23 control). To analyse grooming duration, we constructed a linear mixed effects model with duration (seconds) as a continuous dependent variable, and experimental treatment (pathogen-experience or control) as a discrete explanatory variable. We log-transformed the response variable due to the original model violating assumptions of normality, homoscedasticity, and linearity. To investigate grooming frequency, we constructed a generalised linear mixed effects model with a Poisson error structure and logit-link function. We included an observation-level random intercept effect to account for model overdispersion (Harrison, 2014). To analyse grooming latency, we used a Cox proportional hazards regression model, which compares the time to an event occurring (grooming) across treatments. We also investigated the effect of groomer experience on the number of fungal conidiospores remaining on larvae after the experimental period. Our models included three levels within our fixed effect: larvae groomed by control ants, by UV-pathogen-experienced ants, and larvae not groomed at all. We fitted a generalised linear mixed effects model to this data using a Poisson distribution with a logit-link function. Since this model showed overdispersion, we also included an observation-level random intercept effect (Harrison, 2014), and this model satisfied assumptions of no overdispersion, multicollinearity, or outliers. Lastly, we analysed whether the number of conidiospores remaining after grooming changed during our experimental period (comparing fungal-experienced ants’ first and last exposure periods). We constructed a linear mixed effects regression model, with conidiospore number as the response variable, and experimental time-point as a fixed effect. Since multiple measurements (start and end of experiment) were taken for each ant, we included a random intercept effect for ant identity, and colony of origin. We found this model violated assumptions of linearity, homogeneity of variance, and normality of residuals, however this was fixed by log-transforming the response variable.

### Group experiments

To investigate behaviour in groups, we performed a scan sample of individual behaviour every 4 minutes for 1.5h on the final ‘test day’. To analyse this data, we summed the behaviours for each individual (n=23 time points). Since recordings of allogrooming and venom gland grooming were very low (less than 1% of total observations), we did not include these behaviours in our analysis. We analysed each behaviour individually by constructing a model with experimental treatment (pathogen-experienced or control, for both UV-irradiated and live conidiospores) as a fixed effect, using a generalised linear mixed effects model with a Poisson error structure and logit-link function. In our inactivity model, we removed three observations for ants which showed unusual levels of lethargy and were subsequently outliers (n = 1 from UV-pathogen-experience, live-pathogen-experience, and live control groups). Details of the final models used for analyses are given in Table 1.

**Table 1.** Final statistical models used to analyse group experiment. All models were fitted using the ‘lme4’ package in R, fitting a generalised linear mixed effect model with experimental treatment (UV-pathogen experience, live-pathogen experience, control ants exposed to UV-irradiated conidiospores during the test phase, control ants exposed to live (infectious) conidiospores during the test phase) as a fixed effect. All models were fitted with a Poisson distribution with a logit-link function.

| Dependent variable | Fixed effect | Random intercept effect |
| --- | --- | --- |
| Brood grooming | Experimental treatment | Experimental session<br>Colony of origin<br>Observation |
| Self-grooming | Experimental treatment | Experimental session<br>Colony of origin<br>Observation |
| Brood carrying | Experimental treatment | Experimental session<br>Colony of origin |
| Sitting with the brood | Experimental treatment | Observation |
| Walking | Experimental treatment | Experimental session<br>Observation |
| Inactivity | Experimental treatment | Experimental session<br>Colony of origin<br>Observation |

### Naïve nestmate experiments

To investigate behaviour when groups differed in the workers’ pathogen-experience levels, we formed groups consisting of two pathogen-naïve ants, and a third nestmate that had previously been exposed to either the live or UV-irradiated form of the pathogen or had experienced only the experimental procedure. After introducing pathogen-infected larvae, we conducted scan samples of the behaviour of each ant, summing observations over the 1.5h period (n = 23 timepoints). As we did not differentiate between the two pathogen-naïve ants, we randomly assigned each worker to one of the two possible recorded behaviours before summation (as our focus was on the difference between naive and experienced ants rather than individual ant behaviour). Since recordings of sitting with the brood, carrying the brood, and venom gland grooming were very low (each <1% of total observations), these were not analysed. For each behaviour, we used a Poisson distribution with a logit-link function to model the frequency of occurrence during the scan sample. Our fixed effect in these models (‘condition’) had eight levels representing combinations of pathogen-naïve and pathogen-experienced ants across the four pathogen treatments from the previous group experiment. Details of the final statistical models are given in Table 2.

**Table 2.** Final statistical models used to analyse naïve nestmate experiments. All models were fitted using the ‘lme4’ package in R, fitting a generalised linear mixed effect model with condition (ants from group experiment or naïve nestmates, for each of four pathogen treatments) as a fixed effect. All models were fitted with a Poisson distribution with a logit-link function.

| Dependent variable | Fixed effect | Random intercept effect |
| --- | --- | --- |
| Brood grooming | Condition | Experimental session<br>Colony of origin<br>Observation |
| Self-grooming | Condition | Colony of origin<br>Observation |
| Allogrooming | Condition | Colony of origin |
| Walking | Condition | Colony of origin<br>Observation |
| Inactivity | Condition | Colony of origin<br>Observation |

## RESULTS

### Individual experiments

We investigated the effect of experimental treatment (UV-pathogen-experienced or control) on three aspects of larval grooming behaviour: (i) duration of each grooming event, (ii) frequency, the number of grooming events, and (iii) latency, the time until first instance of grooming. We found no significant effect of treatment on any of our measures of grooming behaviour (Duration: Likelihood ratio test (LR)_df_: *χ²_1_* = 1.49, *P* = 0.22; Frequency: LR: *χ²_1_* = 0.27, *P* = 0.60; Latency: Cox proportional hazards regression: *χ²_1.57_* = 2.57, *P* = 0.20; Figure 5). Similarly, we found no effect of pathogen experience on the number of conidiospores on larvae after grooming (LR: *χ²_2_* = 0.10, *P* = 0.95; Figure 5a), nor on the number of conidiospores remaining after grooming as an ant gained experience during the experiment (LR: *χ²_1_* = 2.03, *P* = 0.15; Figure 5b).

**Figure 5.**
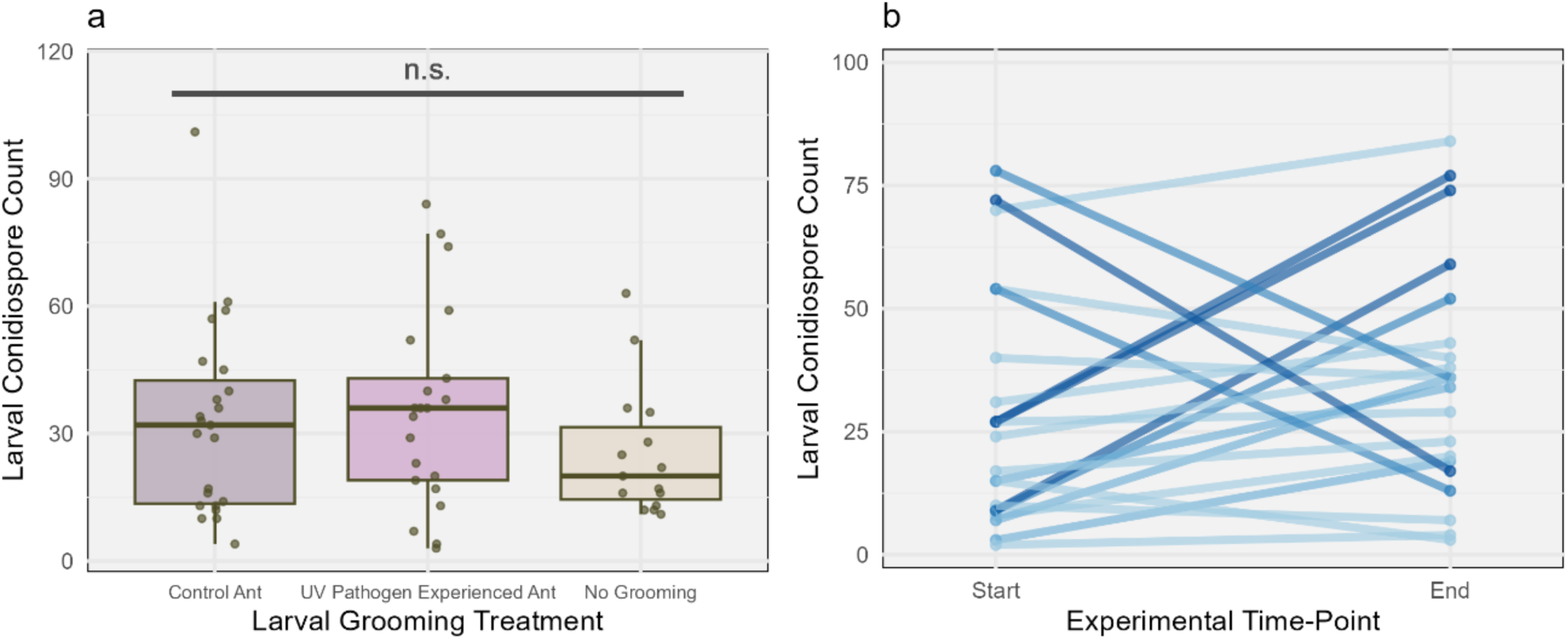
Conidiospore counts during individual experiments. (A) Number of conidiospores remaining on larvae after grooming by control ants (grey), UV-pathogen-experienced ants (purple), and no grooming controls (beige). Boxplot features: centre line = median, box limits = 1^st^ and 3^rd^ quartiles (Q1 to Q3), whiskers = from Q1 to smallest value, at most –1.5 × IQR (inter-quartile range; Q3-Q1), and from Q3 to largest value, at most +1.5 × IQR. All data points displayed, jitter = 0 vertically, 0.1 horizontally. Letters above boxplots depict results of pairwise significance testing between groups, n.s. = non-significant; (B) Number of conidiospores remaining on larvae after grooming by the same ant over the experimental period. Lines coloured by assigning the absolute value of the difference to bins of size 15, and assigning each bin a shade of blue, with larger absolute values coloured by darker shades.

### Group experiments

We found significant effects of experimental treatment on frequency of brood grooming, brood carrying, and sitting with the brood (brood grooming: LR: *χ²_3_* = 11.11, *P* = 0.01; brood carrying: LR: *χ²_3_* = 12.95, *P* = 0.007; sitting with brood: LR: *χ²_3_* = 15.44, *P* = 0.004; Figure 6). Our post-hoc comparisons of brood grooming behaviour found a significant difference between control groups given larvae treated with UV-irradiated conidiospores, and control groups given larvae treated with live (infectious) conidiospores (Tukey post-hoc comparisons: UVcontrol vs. LIVEcontrol *P* = 0.004, all others *P* > 0.05; Figure 6a). Post-hoc analyses of brood carrying behaviour found significant differences between live-pathogen-experience groups and both the experienced and control groups given larvae coated in UV-irradiated conidiospores (Tukey post-hoc comparisons: UVexperienced vs. LIVEexperienced *P* = 0.02, UVcontrol vs. LIVEexperienced *P* = 0.04, all others *P* > 0.05; Figure 6b). Our post-hoc comparisons of sitting with the brood behaviour found a significant difference between pathogen-naïve control groups given larvae treated with UV-irradiated pathogen and all other groups (Tukey post-hoc comparisons: UVcontrol vs. LIVEexperienced *P* = 0.048, UVcontrol vs. UVexperienced *P* = 0.001, UVcontrol vs. LIVEcontrol *P* = 0.008, all others *P* > 0.05; Figure 6c). We found no significant effect of treatment on self-grooming, walking, or inactivity (self-grooming: LR: *χ²_3_* = 6.55, *P* = 0.09; walking: LR: *χ²_3_* = 6.93, *P* = 0.07; inactivity: LR: *χ²_3_* = 5.35, *P* = 0.15; Figure 7).

**Figure 6.**
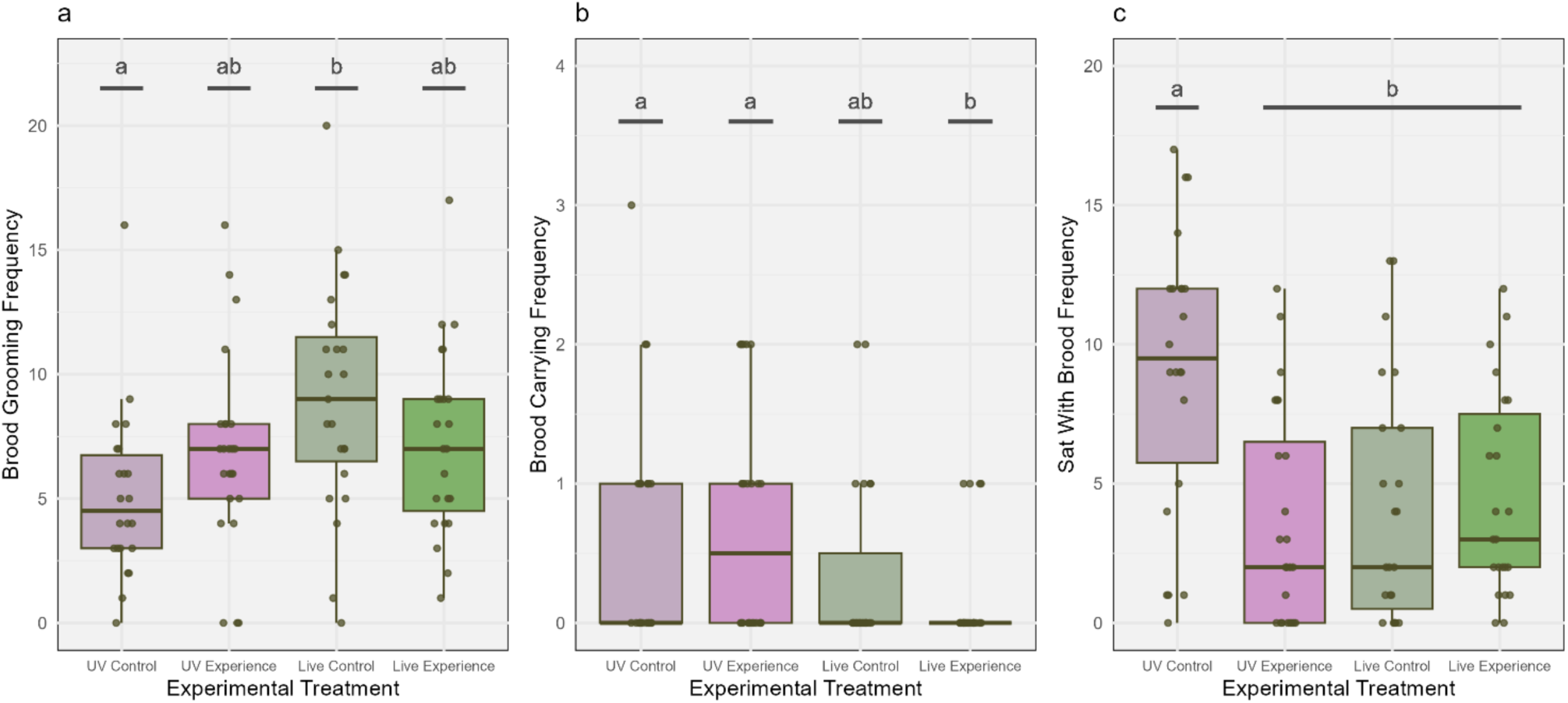
Significant behavioural effects of group pathogen experience. Boxplots show the frequencies of (A) brood grooming, (B) brood carrying, and (C) sitting with brood behaviours observed during the experimental period for control ants given larvae coated in UV-irradiated pathogen (dark purple boxplots), UV-pathogen-experienced ants given larvae coated in UV-irradiated pathogen (light purple boxplots), control ants given larvae coated in live (infectious) pathogen (dark green boxplots), and live-pathogen-experienced ants given larvae coated in live (infectious) pathogen (light green boxplots). Boxplot features: centre line = median, box limits = 1^st^ and 3^rd^ quartiles (Q1 to Q3), whiskers = from Q1 to smallest value, at most –1.5 × IQR (inter-quartile range; Q3-Q1), and from Q3 to largest value, at most +1.5 × IQR. Letters above boxplots depict results of pairwise significance testing between groups. All data points displayed, jitter = 0 vertically, 0.1 horizontally.

**Figure 7.**
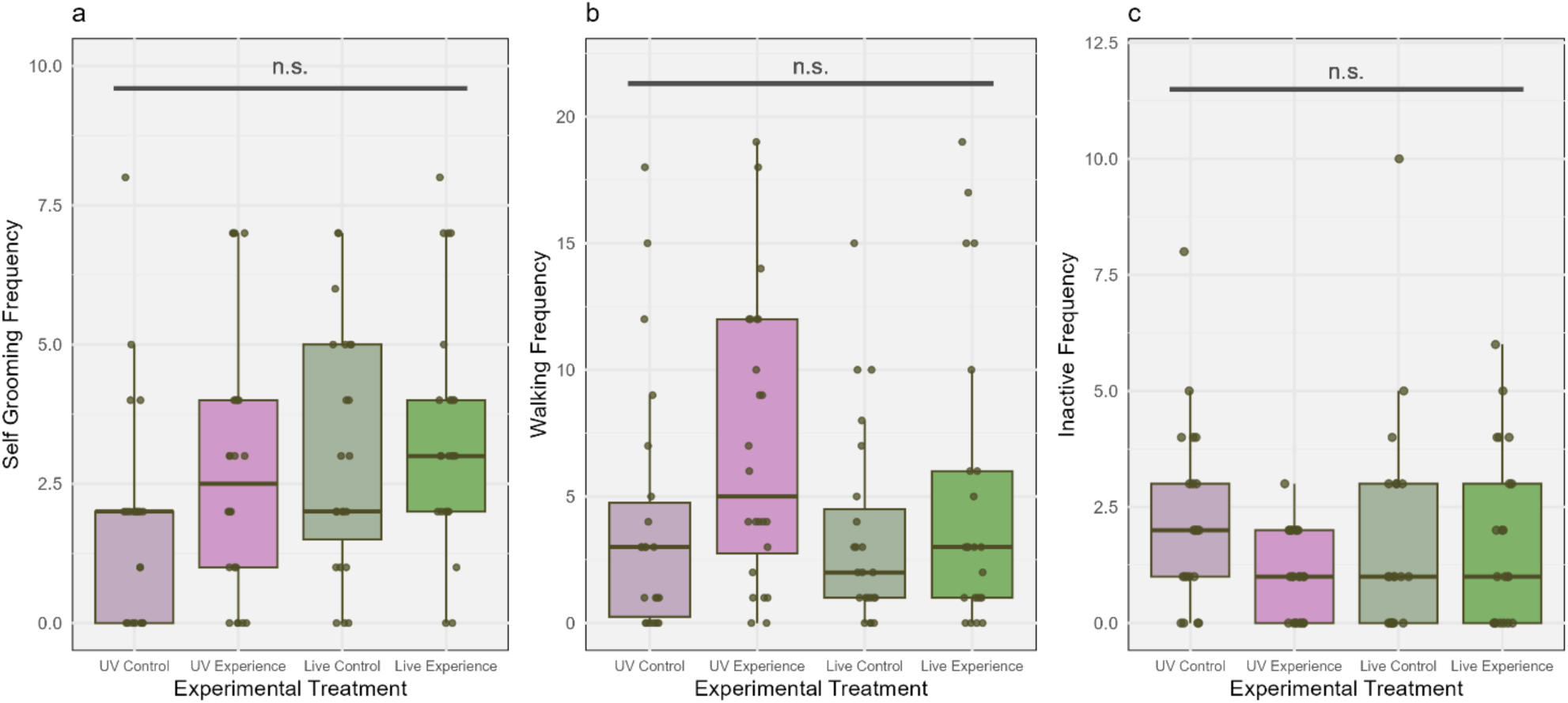
Non-significant behavioural effects of group pathogen experience. Boxplots show the frequencies of (**A**) self-grooming, (**B**) walking, and (**C**) inactive behaviours observed during the experimental period for control ants given larvae coated in UV-irradiated pathogen (dark purple boxplots), UV-pathogen-experienced ants given larvae coated in UV-irradiated pathogen (light purple boxplots), control ants given larvae coated in live (infectious) pathogen (dark green boxplots), and live-pathogen-experienced ants given larvae coated in live (infectious) pathogen (light green boxplots). Boxplot features: centre line = median, box limits = 1^st^ and 3^rd^ quartiles (Q1 to Q3), whiskers = from Q1 to smallest value, at most –1.5 × IQR (inter-quartile range; Q3-Q1), and from Q3 to largest value, at most +1.5 × IQR. Letters above boxplots depict results of pairwise significance testing between groups, n.s. = non-significant. All data points displayed, jitter = 0 vertically, 0.1 horizontally

### Naive nestmate experiments

We found a significant effect of condition on allogrooming and walking (Allogrooming: LR: *χ²_7_* = 14.16, *P* = 0.048; Walking: LR: *χ²_7_* = 15.23, *P* = 0.048; Figure 8). However, for both behaviours, our post-hoc analyses found no significant pairwise interactions. We found no significant effect of condition on brood grooming, self-grooming, or inactivity (Brood grooming: LR: *χ²_7_* = 12.47, *P* = 0.09; Self-grooming: LR: *χ²_7_* =11.78, *P* = 0.11; Inactivity: LR: *χ²_7_* =12.46, *P* = 0.09; Figure 10).

**Figure 8.**
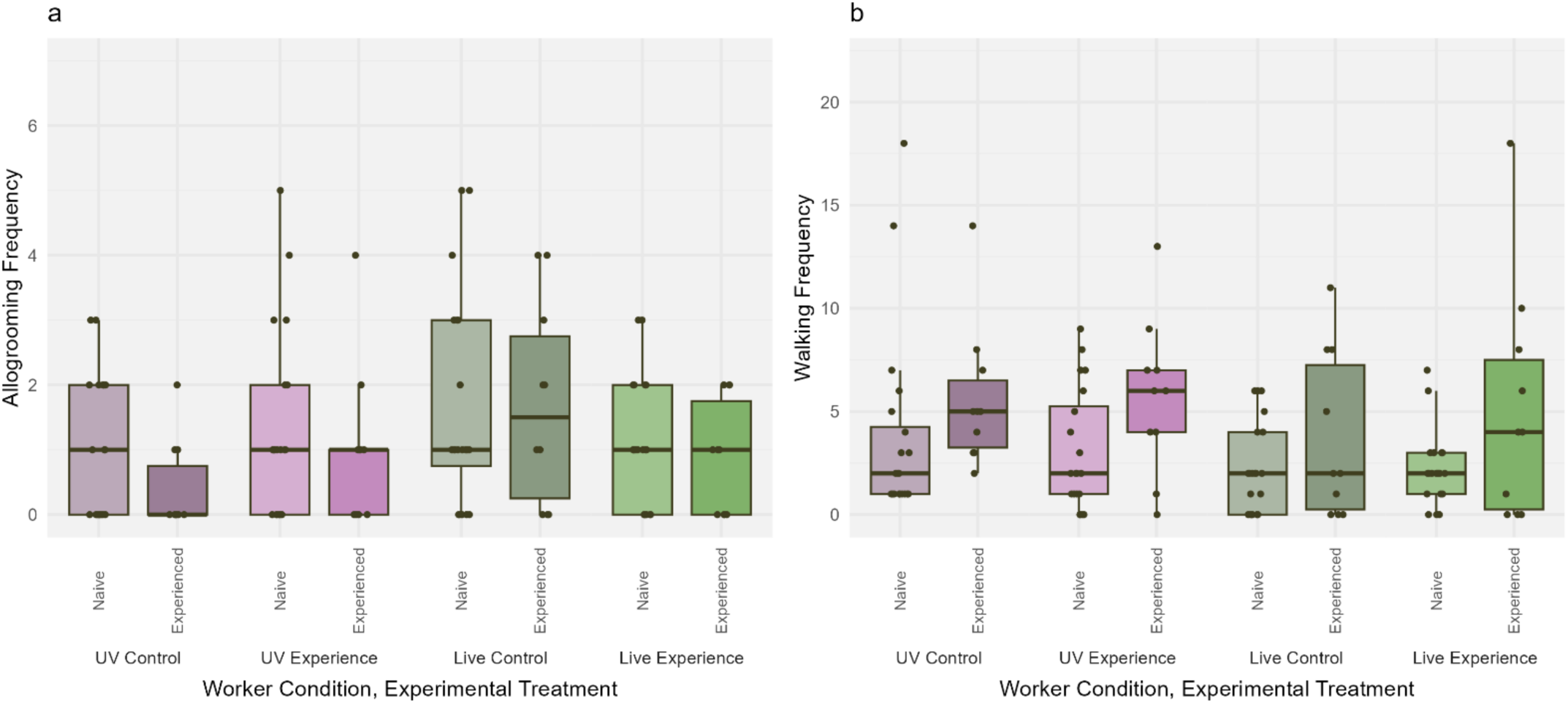
Significant behavioural effects in naïve nestmate experiments. Boxplots show the frequencies of (**a**) allogrooming, and (**b**) walking behaviours observed during the experimental period for pathogen-naïve nestmates (‘Nestmate’; darker colour boxplots) and ants from prior experiments (‘Experimental’; lighter colour boxplots). These ants were housed in groups of three where the experimental ant had no prior pathogen experience and groups were given a larva coated in UV-irradiated pathogen (dark purple boxplots), the experimental ant was UV-pathogen-experienced and groups were given a larva coated in UV-irradiated pathogen (light purple boxplots), the experimental ant had no prior pathogen experience and groups were given a larva coated in live pathogen (dark green boxplots), or the experimental ant was live-pathogen-experienced and groups were given a larva coated in live (infectious) pathogen (light green boxplots). Boxplot features: centre line = median, box limits = 1^st^ and 3^rd^ quartiles (Q1 to Q3), whiskers = from Q1 to smallest value, at most –1.5 × IQR (inter-quartile range; Q3-Q1), and from Q3 to largest value, at most +1.5 × IQR. All data points displayed, jitter = 0 vertically, 0.1 horizontally.

**Figure 9.**
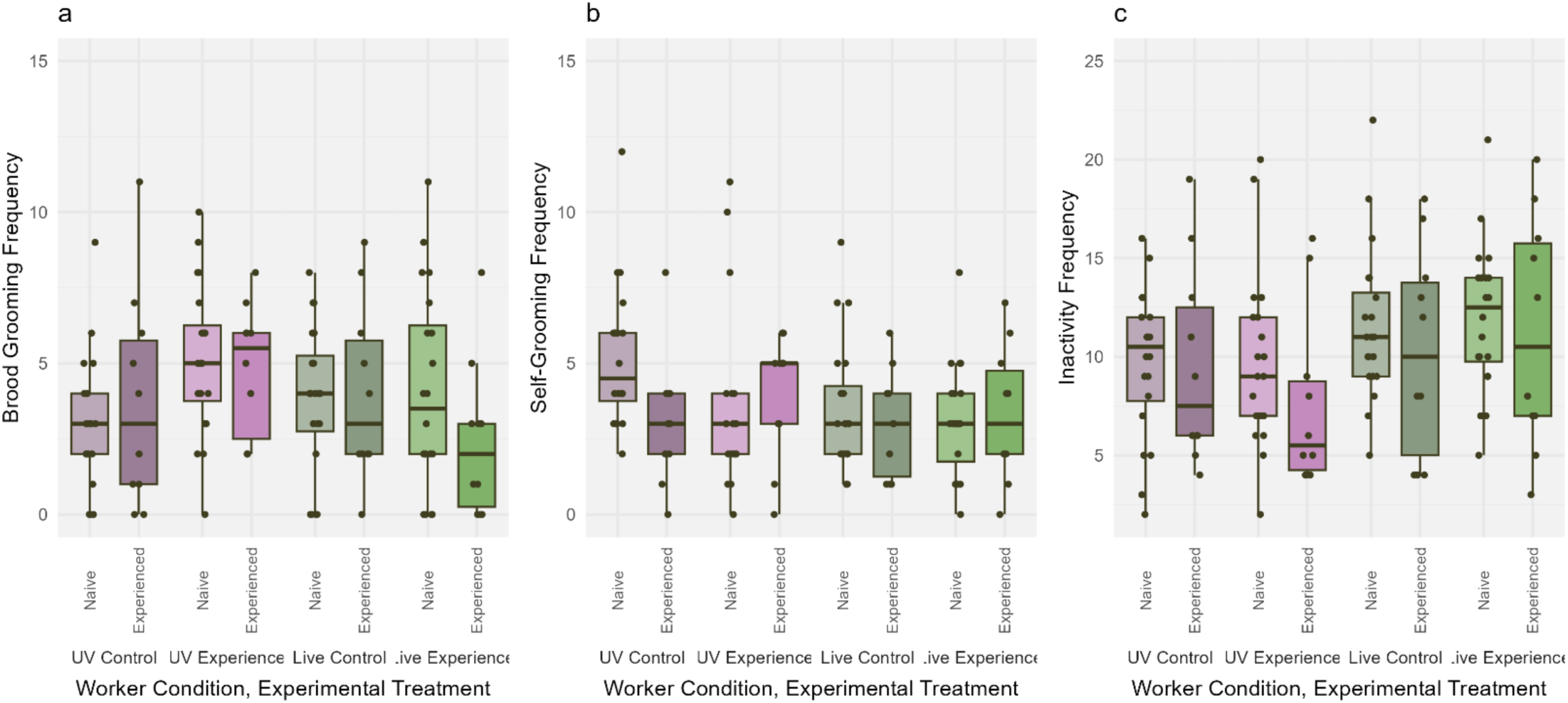
Non-significant behavioural effects in naïve nestmate experiments. Boxplots show the frequencies of (A) brood grooming, (B) self-grooming, and (C) inactivity behaviours observed during the experimental period for pathogen-naïve nestmates (‘Nestmate’; darker colour boxplots) and ants from prior experiments (‘Experimental’; lighter colour boxplots). These ants were housed in groups of three where the experimental ant had no prior pathogen experience and groups were given a larva coated in UV-irradiated pathogen (dark purple boxplots), the experimental ant was UV-pathogen-experienced and groups were given a larva coated in UV-irradiated pathogen (light purple boxplots), the experimental ant had no prior pathogen experience and groups were given a larva coated in live pathogen (dark green boxplots), or the experimental ant was live-pathogen-experienced and groups were given a larva coated in live (infectious) pathogen (light green boxplots). Boxplot features: centre line = median, box limits = 1^st^ and 3^rd^ quartiles (Q1 to Q3), whiskers = from Q1 to smallest value, at most –1.5 × IQR (inter-quartile range; Q3-Q1), and from Q3 to largest value, at most +1.5 × IQR. All data points displayed, jitter = 0 vertically, 0.1 horizontally.

## DISCUSSION

In this study, we examined whether prior experience with a fungal pathogen, both individually and in groups, affects ant social immunity behaviour, and examined the effect of prior experience when groups differ in their pathogen experience levels. We found that (i) in our individual experiments, prior pathogen experience had no effect on brood grooming behaviour relative to no-pathogen-experience controls (Figure 4); (ii) similarly, groomer experience level had no effect on the number of fungal conidiospores removed from larvae (Figure 5a) or by the same ant during the experimental program (Figure 5b); (iii) when building pathogen experience in groups, brood grooming, brood carrying, and sitting with the brood behaviours differed between treatments, but not between our pathogen-experience and control groups (Figure 6); and (iv) when pathogen-experienced ants were placed with naïve nestmates, this had some effect on allogrooming and walking behaviour (Figure 8), yet no two groups of ants were significantly different from one another.

In contrast to previous results (Goes et al., 2023; Walker & Hughes, 2009), our study finds no evidence that prior pathogen experience affects social immunity behaviour when ants are secondarily presented with a pathogen. This could have resulted from the use of a smaller group size in our experiments; while the largest group size in our experiments was only five ants, previous effects of pathogen experience were demonstrated in groups of 21 workers (Walker & Hughes, 2009) or whole colonies (Goes et al., 2023). Such differences in group size may affect the development of experience-induced effects. For example, our experiments may more closely resemble pathogen encounter outside the nest environment, where individuals often avoid infected stimuli (Mehdiabadi & Gilbert, 2002; Westhus et al., 2014), or they may have been impacted by social isolation from the colony, which can affect brood care behaviour (Scharf et al., 2021). Additionally, previous reports of experiential effects were found over longer time periods than those used in our experiments. Whereas Walker & Hughes (2009) exposed groups to pathogen-treated ants for 28 days and Goes and colleagues (2023) found significant increases in hygienic behaviour after 1 and 2 months, our experiments only lasted six days. A shorter experimental period may, therefore, have not met a threshold level of pathogen exposure that invokes experience-induced effects. Further, while our experiments found no effect of pathogen infection in producing increases in grooming behaviour (Figures 7 and 8, green boxplots), the longer time periods of previous experiments may have resulted in ants building higher pathogen loads, as well as encountering additional infection-related stimuli (e.g., sporulating corpses), which may be necessary for triggering experience-induced responses.

The use of Triton-X100 in our experiments may also have confounded results. This detergent was required to infect ants with *Metarhizium* conidiospores due to their hydrophobicity, and is widely used in similar social immunity studies (e.g. Walker & Hughes, 2009; Westhus et al., 2014). However, it has previously been suggested that this detergent may act as an irritating compound to ants (Westhus et al., 2014). In the context of our experiments, therefore, comparing pathogen-experience to that with an irritant may have biologically different interpretations to comparisons with experience of uncontaminated larvae. Indeed, we did not observe any differences in social immunity behaviour between the pathogen and control treatments in most cases, and alternative mechanism to introduce pathogens in social immunity experiments should possibly be explored.

Conversely, we may not have detected a ‘social immune memory’ effect simply because the ants used in this experiment do not demonstrate this behaviour. Instead, sufficient protection against pathogens may be achieved by other social immunity responses. For example, it has been demonstrated that ants show elevated grooming after experiencing disease, regardless of pathogen presence (Morelos-Juárez et al., 2010; Reber et al., 2011; Westhus et al., 2014). Since parasite invasions often come in ‘waves’ (Schmid-Hempel, 2013a), this prophylactic upregulation in anti-pathogen behaviour may achieve many of the infection-reducing benefits of utilising prior experience, while potentially having lower metabolic costs (Burns et al., 2011). However, without integrating prior experience, this behaviour could result in erroneous grooming, which can also be costly to colonies (Reber et al., 2011). The existence of an experienced-induced effect may also be species-specific. In contrast to ant species more often used to investigate social immunity behaviours (e.g., *Lasius*, Pull et al., 2018), *Temnothorax* ants frequently relocate nests, and so perhaps abandon their nest and infected/contaminated brood instead of investing into costly social immunity measures. Ants may, therefore, demonstrate species-specific biases towards different social immunity strategies, such as grooming versus nest relocation. Indeed, species may differ significantly in their responses to pathogen exposure, given their differences in life histories and ecologies, and the differing methods and intensities of infection that may result from this. We therefore call for more studies of social immunity responses outside of established “model” species to investigate these potential differences further.

In our group experiments, we found that ants who had experienced the live pathogen spent less time carrying the brood than both pathogen-experienced and pathogen-naive groups that were given larvae with UV-irradiated conidiospores (Figure 6b). Live-pathogen-experienced ants had likely contracted infections during the experimental period (Vestergaard et al., 1999), and our results are therefore in agreement with previous findings that infected ants spend less time in the brood chamber, potentially limiting infections transmission within colonies (Stroeymeyt et al., 2014; Ugelvig & Cremer, 2007).

### Conclusions & Further Directions

This study reports no significant effect of pathogen experience on ant behaviour during pathogen reencounter. This is nonetheless an interesting result and may suggest that experience-induced effects can be flexibly employed, depending on the context in which experience occurs, or conversely, may represent species-specific differences in pathogen response behaviours. Further experimentation could, therefore, explore whether this behaviour is demonstrated when groups are larger, and experience is built over a longer time, or if different conidiospore application methods result in stronger differences between pathogen-treatment and control groups. Similarly, it would be beneficial to investigate the impact of erroneous brood grooming, to determine the potential benefits arising from experience-induced effects, and to investigate potential interspecies differences in pathogen responses (e.g., brood grooming vs nest reallocation). In contrast to organismal immune memory, social immune memory thus appears to be a more flexibly demonstrated trait, with alternative pathogen-response mechanisms perhaps operating between different scenarios, species, or indeed across biological scales.

## ACKNOWLEDGEMENTS

We thank Natalie Stroeymeyt for advice regarding ant painting techniques, and Greg Pask and Ben Morris for guidance setting up the Raspberry Pi filming system.

